# Local genetic neighbourhoods but resilient gene flow across anthropogenic landscapes in the red campion (*Silene dioica*)

**DOI:** 10.64898/2026.06.05.730353

**Authors:** Camille Jolivel, Jean-François Arnaud, Estelle Barbot, Cécile Godé, Nina Joffard, Isabelle De Cauwer

## Abstract

**Background and Aims:** In the plant kingdom, gene flow occurs through pollen and seed dispersal, shaping both within-population spatial genetic structure and among-population genetic differentiation. Anthropogenic land-use change can affect levels of gene flow by reducing pollinator abundance and altering pollen and seed dispersal pathways. Yet, how environmental features shape gene flow events in common herbaceous plants – a fundamental building block of many ecosystems – remains poorly understood. We address this gap by investigating how environmental context influences population genetic structure at regional and local scales in the red campion (*Silene dioica*).

**Methods:** By sampling 1,005 individuals from 29 populations across habitats ranging from semi-natural to strongly human-altered, we assessed whether population size and landscape composition influenced within-population genetic diversity and population genetic differentiation. At the local scale, we examined whether landscape composition affected fine-scale spatial genetic structure and pollen dispersal distances in a subset of six populations representing the two extremes of an anthropogenic gradient.

**Key Results:** No effect of either demographic or landscape factors was found on levels of genetic diversity. We detected moderate levels of genetic differentiation among populations that matched an isolation-by-distance pattern, with high levels of population admixture, while landscape composition did not explain variation in population genetic differentiation. At the local scale, five of the six studied populations exhibited significant spatial genetic structuring, indicating distinct neighbourhoods at spatial scales less than 10 m. Paternity analyses based on 4,800 offspring further revealed predominantly short-distance pollen dispersal together with substantial immigration from external sources. Neither fine-scale genetic structure nor pollen dispersal distances differed between habitat types.

**Conclusions:** Altogether, our results demonstrate a multimodal pattern of gene flow and a remarkable resilience of this common herbaceous species to anthropogenic habitat change, as substantial genetic connectivity is maintained despite insect-mediated pollen dispersal and gravity-driven seed dispersal.

## INTRODUCTION

Anthropogenic activities, notably urban and agricultural expansion, have profoundly altered species distributions and demographic dynamics by transforming continuous habitats into fragmented landscapes (Haddad *et al*. 2015; Newbold *et al*. 2015). Such environmental changes typically reduce population sizes and increase spatial isolation among populations, ultimately shaping patterns of intraspecific genetic diversity (Slatkin 1985; Lowe *et al*. 2004; Ellegren and Galtier 2016). By decreasing effective population sizes, fragmentation indeed enhances levels of inbreeding and genetic drift within populations and can lead to rapid erosion of genetic variation, as documented in both plant and animal species (Young *et al*. 1996; Van Rossum *et al*. 2004; Honnay and Jacquemyn 2007; Haag *et al*. 2010; Frankham *et al*. 2017; Perrin *et al*. 2021; Allendorf *et al*. 2022). Simultaneously, reduced connectivity among habitat patches limits gene flow, thereby increasing genetic differentiation among populations (Slatkin 1985; Lowe *et al*. 2004; Frankham *et al*. 2017). These genetic consequences can arise over relatively short timescales, particularly in species with short generation times. Agricultural landscapes, which are among the environments most affected by human activities, have experienced intensified management since the mid-20th century (Matson *et al*. 1997; Benton *et al*. 2003). This period spans many generations for short-lived herbaceous species, resulting in detectable signatures in spatial genetic structure (SGS) in some taxa (e.g., Berge *et al*. 1998; Emel *et al*. 2021; Guiller *et al*. 2023; Huang *et al*. 2025; Lamperty *et al*. 2025), including several common species (e.g., Van Rossum *et al*. 2004; Honnay and Jacquemyn 2007).

Anthropogenic impacts on genetic structure are not uniform across the plant kingdom: species traits, particularly dispersal ability – including the two main pathways of plant gene flow, namely pollen and seed dispersal – can determine the magnitude of these effects and influence population vulnerability to anthropogenic habitat change. The relative contribution of the pollen and seed dispersal to overall gene flow can further modulate the magnitude of anthropogenic impacts, with pollen-mediated gene flow often playing a key role in connecting subdivided populations (Ennos 1994; Berge *et al*. 1998; Petit *et al*. 2005; Breed *et al*. 2015; Favre-Bac *et al*. 2016; Guiller *et al*. 2023). Compared with wind-pollinated species, animal-pollinated plants are expected to be more strongly impacted by land-use change, since they rely on pollen vectors that are themselves affected by anthropization. In particular, agricultural intensification negatively affects both pollinator abundance and diversity through the intensive use of pesticides and herbicides, as well as through reductions in floral resources and nesting sites (Winfree *et al*. 2009; Woodard and Jha 2017), while decreasing landscape connectivity (Aguilar *et al*. 2008; Aavik *et al*. 2013). Pollinator decline can in turn constrain seed production, reducing plant population size (Biesmeijer *et al*. 2006) and potentially increasing genetic differentiation among populations through genetic drift (Sullivan *et al*. 2019). Land-use change can also alter pollinator assemblages, with potential consequences for pollen flow and plant SGS, since pollinator functional traits such as body size and mobility are key determinants of pollen flow. For instance, plant species pollinated by small insects often exhibit higher levels of SGS than those pollinated by larger-bodied insects (Gamba and Muchhala 2020), likely because larger pollinators forage over longer distances and promote more extensive pollen-mediated gene flow (e.g., Zurbuchen *et al*. 2010). In landscape mosaics composed of patches of anthropogenic and semi-natural habitats, pollinator assemblages may vary markedly across space (e.g., Herrera 1988; Gómez *et al*. 2008), which, in generalist plant species (i.e., those pollinated by multiple functional groups of insects), may translate into spatially heterogeneous patterns of pollen flow and, consequently, genetic differentiation among populations.

As with pollen dispersal, there is a diversity of seed dispersal modes – including gravity, ballistic, wind, and animal-mediated dispersal – which can influence dispersal distances (Hamrick and Godt 1996; Petit *et al*. 2005; Choo *et al*. 2012; De Cauwer *et al*. 2012; Favre-Bac *et al*. 2016; Latron *et al*. 2020). For example, gravity-dispersed seeds typically travel shorter distances than seeds dispersed by other modes (Thomson *et al*. 2011). Yet, comparative studies have not consistently detected differences in SGS among species exhibiting distinct dispersal modes (Hamrick and Godt 1996; Duminil *et al*. 2007; Gamba and Muchhala 2020), possibly due to the stochastic nature of seed dispersal and because seed-mediated gene flow typically represents only a limited fraction of overall gene flow among populations (Petit *et al*. 2005). Beyond facilitating population connectivity, seed dispersal is essential for the colonization of suitable habitats, thereby promoting the successful establishment of new populations (McCauley *et al*. 1995; Cain *et al*. 2000; Fievet *et al*. 2007; Dellafiore *et al*. 2010; Latron *et al*. 2020). This may be particularly important in unstable or transient habitats, such as those found in urban or agricultural landscapes, where suitable patches are frequently created, modified, or destroyed (McCauley *et al*. 1995; Favre-Bac *et al*. 2016; Guiller *et al*. 2023; Hardion *et al*. 2026). Importantly, anthropogenic pressures may alter patterns of seed dispersal. In particular, land-use change may disrupt animal-mediated dispersal by altering the movement of seed vectors (Fricke *et al*. 2025). Conversely, agricultural activities may occasionally facilitate long-distance dispersal via machinery, promoting gene flow in human-disturbed landscapes (Strykstra *et al*. 1997; Klinger *et al*. 2021). Overall, anthropogenic habitat change is expected to strongly affect spatial patterns of pollen and seed flow, although its effects on SGS remain difficult to predict and likely depend on species traits (Aguilar *et al*. 2008; Favre-Bac *et al*. 2016; Auffret *et al*. 2017).

Although regional patterns of genetic connectivity are largely determined by gene flow among populations, dispersal mechanisms can also strongly influence the spatial distribution of genetic diversity at much finer spatial scales (Heywood 1991; Epperson 1995). Within populations, local SGS is largely determined by pollen movement among neighbouring individuals and short-distance seed dispersal (Ingvarsson and Giles 1999; Vekemans and Hardy 2004; Van Rossum and Triest 2007; Choo *et al*. 2012; De Cauwer *et al*. 2012; Miguel-Peñaloza *et al*. 2023). Even in plant species pollinated by highly mobile insects – potentially capable of connecting distant populations – pollen movement within populations is often spatially restricted (Heywood 1991; Hardy *et al*. 2004; Van Rossum and Triest 2007; Miguel-Peñaloza *et al*. 2023). Once pollinators reach a patch of plants, they typically forage sequentially among nearby individuals, optimizing energy intake while minimizing movement costs (Schmitt 1980). Such short-distance foraging behavior can generate fine-scale SGS within populations by creating local isolation-by-distance patterns, whereby genetic relatedness declines with increasing distance among individuals (Wright 1943; Epperson 1995; Ingvarsson and Giles 1999; Vekemans and Hardy 2004; Isagi *et al*. 2007; Van Rossum and Triest 2007; Lamperty *et al*. 2025). Fine-scale SGS tends to be particularly pronounced in species combining self-fertilization, limited seed dispersal and herbaceous growth form (Heywood 1991; Loiselle *et al*. 1995; Vekemans and Hardy 2004; Latron *et al*. 2020). In fragmented habitats, these trait-dependent patterns may be further amplified. Habitat fragmentation can increase selfing rates through reduced local plant density and/or altered pollinator behavior in sparse patches (Setsuko *et al*. 2013; Breed *et al*. 2015), thereby strengthening fine-scale SGS (Raabová *et al*. 2015). In contrast, in strictly outcrossing species, such as those exhibiting self-incompatibility or dioecy, reduced plant density may weaken fine-scale SGS by increasing within-population pollination distances (e.g., Isagi *et al*. 2007). Consistent with this conceptual framework, empirical studies report contrasting outcomes, with habitat fragmentation shown to either strengthen (Van Rossum and Triest 2007; De-Lucas *et al*. 2009) or weaken (Born *et al*. 2008) fine-scale SGS, whereas other studies have found little or no detectable effect (Smith *et al*. 2018). A recent meta-analysis further shows that available data remain too scarce, particularly for herbaceous species, to draw general conclusions about the effect of anthropization on fine-scale SGS (Miguel-Peñaloza *et al*. 2023). In particular, while pollen dispersal distance in herbaceous species is often spatially restricted (e.g., DiLeo *et al*. 2018; Barbot *et al*. 2022), the extent to which anthropogenic habitat change alters these patterns remains unknown, underscoring the need to conduct paternity analyses in populations located in contrasting (i.e., semi-natural vs. human-altered) habitats.

In this study, we investigated spatial patterns of neutral genetic variation at both regional and local scales in the common dioecious herbaceous species *Silene dioica*. Spatial genetic structure integrates the cumulative effects of gene flow and drift over time and can therefore reveal the long-term impacts of anthropogenic land-use change. To complement this historical perspective, we used paternity analyses to quantify contemporary pollen dispersal at local scales, thereby providing a snapshot of ongoing gene flow. Although typically described as a forest-associated plant, this species is capable of colonizing a wide range of habitats, including field edges, roadsides, and other anthropogenic environments, provided that conditions remain partially shaded and soils moist (Giles and Goudet 1997). We sampled 29 populations spanning natural variation in population size and habitat context in a landscape mosaic composed of anthropogenic habitats and forest patches in northern France, and genotyped individuals at microsatellite loci. Our objectives were to address the following questions:

1. At the regional scale, do population size and landscape composition influence within-population levels of genetic diversity? We predicted that large populations would harbour higher levels of genetic diversity than small ones, and that populations surrounded by forest and semi-natural habitats would be more genetically diverse than those embedded in predominantly agricultural matrices, where genetic drift may be stronger.
2. At the regional scale, how genetically differentiated are populations across the landscape? Given that the red campion is an insect-pollinated species with gravity-dispersed seeds, we expected substantial genetic differentiation among populations and clear patterns of isolation-by-distance (IBD). We further predicted that populations embedded in predominantly agricultural matrices would be more differentiated than those located in forested or semi-natural landscapes, due to reduced habitat quality and anthropogenic barriers to gene flow.
3. At the local scale, how does habitat influence fine-scale SGS and contemporary pollen dispersal? Focusing on six populations representing contrasting environments (three populations in forest habitats and three in anthropogenic habitats dominated by cultivated fields), we quantified fine-scale SGS and estimated pollen dispersal distances using progeny genotyping and paternity analyses.

By integrating patterns of genetic structure across broad and fine spatial scales with direct estimates of contemporary pollen dispersal, this study provides a multiscale perspective on how landscape configuration shapes patterns of gene flow.

## MATERIALS AND METHODS

### STUDY SPECIES

The red campion (*Silene dioica* (L) Clairv., Caryophyllaceae) is a herbaceous, perennial, and dioecious angiosperm widely distributed in central and northern Europe, often inhabiting semi-natural or anthropogenic habitats (Kay *et al*. 1984; Giles and Goudet 1997). Its lifespan ranges from 5 to 10 years (Giles *et al*. 1998; Goulson 2009), and reproduction is possible from the first year onward. It has a generalist pollination system, primarily mediated by two major taxonomic groups: Hymenoptera, including large insects such as bumblebees and smaller ones such as halictid bees, and Diptera, mainly hoverflies (Kay *et al*. 1984; Barbot *et al*. 2022). Seeds are dispersed by gravity, with no specialized structures facilitating dispersal, and typically germinate the following spring (Giles and Goudet 1997). Seeds can remain dormant, enabling the establishment of a seed bank (Matlack 1987), even though the longevity of the seed bank remains unknown.

### SAMPLING DESIGN AND HABITAT CHARACTERIZATION

Leaf samples from 1,005 individuals were collected across 29 natural populations from northern France and Belgium, each defined as a discrete cluster of individuals separated by at least 100 m from other neighbouring clusters of conspecific individuals. Populations were separated by a mean distance of 30 km (range: 0.27–72.14 km, Figure 1). The species is common and abundant in the study region, so our sampling was not exhaustive but aimed to capture a representative subset of populations in the region. Land-use in the area surrounding each population was quantified using the QGIS software (version 3.30.0) with the GroupStats plugin (version 2.2.7). Land-use was assessed within a one-kilometer radius around each population, corresponding with the typical foraging range of bumblebees (Knight *et al*. 2005), one of the most common pollinators of *S. dioica* (Kay *et al*. 1984; Barbot *et al*. 2022). Four mapping data sources were used depending on the location of the population: the ARCH project, the OS Picardie project, the COSW project, and the THEIA project. Different land-uses were then grouped into four broad habitat types: forest habitats (coniferous and deciduous forests), semi-natural habitats (meadows and pastures), crop habitats (fields and orchards), and urbanized habitats (cities and industrial sites) (Supplementary Table S1). We also measured the distance to the nearest wooded area, considering only forest fragments larger than 300 m². A Principal Component Analysis (PCA) was then performed on habitat characteristics to identify which variables to include in the statistical analyses presented below (Supplementary Fig. S1). PC1, which explained 49.9% of the total variance, indicated that forest habitat surface was negatively correlated with distance to forest, crop habitat surface, and semi-natural habitat surface. PC2, which explained 22.1% of the variance, was primarily associated with urban habitat surface. Forest habitat surface (ranging from 0% to 98%) and urban habitat surface (ranging from 0% to 53%), which best represented PC1 and PC2 respectively, were therefore retained as explanatory variables in the subsequent statistical analyses.

**Figure 1.**
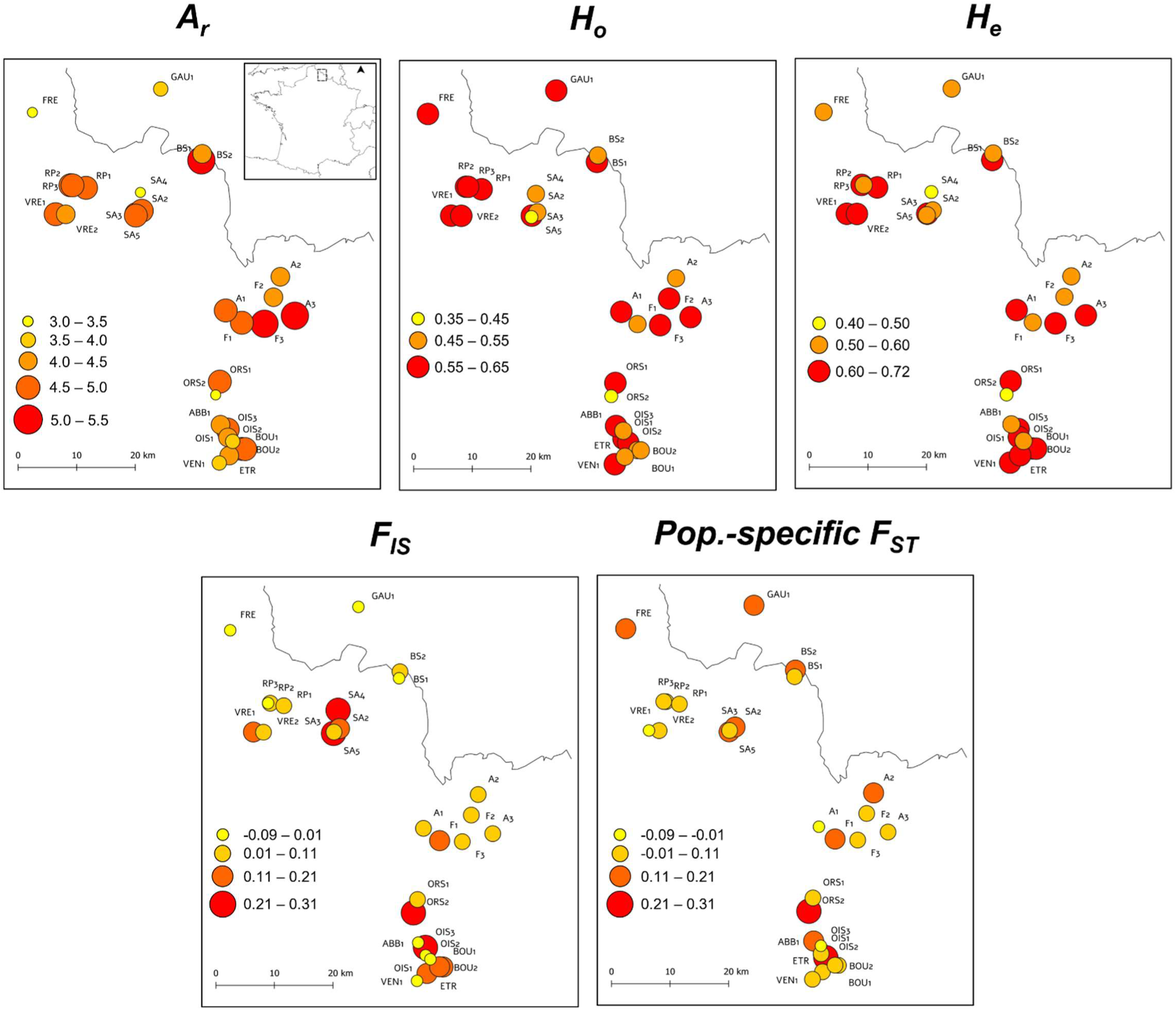
Sampling locations and spatial patterns of population genetic features in red campion (*S. dioica*) populations sampled in northern France and Belgium. The grey line corresponds to the border between countries. *A_r_*: allelic richness; *H_o_*: observed heterozygosity; *H_e_*: gene diversity sensu Nei (1978), *F_IS_*: intra-population fixation index; Pop.-specific *F_ST_*: population-specific *F_ST_* as defined by Weir and Goudet (2017).

To investigate the SGS both among and within populations, we implemented two complementary sampling strategies. Between 2018 and 2019, we carried out broad regional sampling by randomly sampling 20 individuals (when available, mean ± SE = 17.61 ± 0.70 individuals) from 23 natural populations (Supplementary Table S2). This regional sampling was designed to represent the diversity of habitats where the species occurs. In 2023, we further conducted a more intensive sampling within six carefully selected populations with comparable surface areas, representing contrasting habitat types: three forest-dominated populations (labelled “F”) with at least 90% of forest and semi-natural cover in the surrounding landscape, and three anthropogenic populations (labelled “A”) characterized by more than 40% of agricultural and urbanized land in the surrounding landscape. In each of these six populations, 50 males and 50 females were randomly sampled to investigate fine-scale SGS. For all 29 populations, census size was estimated by counting the total number of flowering individuals and ranged from 7 to 2,000 individuals (mean = 234.79, Table S2).

Additional manipulations were carried out in the six intensively sampled populations to standardize demographic conditions and reduce the number of pollen donors, thereby enabling paternity analyses. Prior to peak flowering in 2023, all inflorescences of non-sampled individuals within these populations were removed to ensure comparable population sizes and densities across study sites (50 flowering males and 50 flowering females per population). Paternity analyses were performed on a subsample of the seeds produced during a ten-day window at flowering peak. For each female (50 per population, 300 in total), eight random fruits were harvested at maturity and two seeds per fruit were genotyped, resulting in 16 offspring per female for a total of 4,800 offspring genotyped across all populations.

### MOLECULAR GENOTYPING

Total genomic DNA was extracted from 1,005 adult individuals and 4,800 offspring using the NucleoMag 96 Plant Kit (Macherey-Nagel, Oesingen, Switzerland) and an automated DNA purification system (KingFisher®, Thermo Fisher Scientific, Waltham, USA). DNA extraction protocols for seeds are described in Joffard *et al*. (2025). PCR amplifications were performed at five nuclear microsatellites following Barbot *et al*. (2022). Electrophoresis of PCR products was performed using an automated capillary sequencer (3130 Genetic Analyzer, Applied Biosystems, USA). Multilocus genotypes were scored using GeneMapper (Applied Biosystems, USA). Individuals with ambiguous genotypes were subjected to a second PCR to confirm allele calls. Missing genotypes represented less than 1% of the dataset.

### ANALYSES

#### 1. LEVELS OF GENETIC DIVERSITY AT REGIONAL SCALE

Four measures of genetic diversity were estimated within each population: the allelic richness (*A_r_,* the number of alleles standardized to the size of the smallest sampled population, N = 7, and averaged across the five loci), the multilocus intra-population fixation index (*F_IS_*, a measure of deviation from panmixia), the mean observed heterozygosity (*H_o_*) and the gene diversity (*H_e_*), sensu Nei (1978). These indices were calculated using FSTAT (version 2.9.4, Goudet 2003). The statistical significance of *F_IS_* values was assessed through 1,000 random permutations of alleles among individuals within populations.

The relationships between *A_r_*, *F_IS_*, *H_o_*, *H_e_* and habitat characteristics were tested using linear regressions with habitat characteristics (forest and urban habitat surfaces within a one-kilometer radius around each population) and census population size as explanatory variables. All explanatory variables were centered and scaled (mean = 0, SD = 1) prior to analysis, so that model coefficients represent the effect of a one-standard-deviation change around the mean. Variance inflation factors (VIF < 1.2) indicated minimal multicollinearity among explanatory variables (habitat characteristics and census population size).

For the six populations sampled in 2023, the observed levels of *A_r_*, *F_IS_*, *H_o_* and *H_e_* were compared between habitat types (forest vs. anthropogenic habitats). The statistical significance of differences between habitat types was assessed using 1,000 permutations of individuals among populations and habitat types using FSTAT.

#### 2. LEVELS OF GENETIC DIFFERENTIATION AMONG POPULATIONS AND SPATIAL GENETIC STRUCTURE AT REGIONAL SCALE

We investigated SGS at the regional scale, excluding one population where fewer than 10 individuals could be sampled (see Table S2). First, genetic differentiation across the whole dataset was estimated by the mean multilocus *F_ST_* (Weir and Cockerham 1984) across all populations, using FSTAT. Statistical significance was evaluated using 1,000 random permutations of multilocus genotypes among populations. Second, pairwise *F_ST_* values for all population pairs were computed and their statistical significance was tested using the same permutation procedure with a Bonferroni correction for multiple tests. We then tested the correlation between pairwise *F_ST_* values and geographic distances using Mantel tests (1,000 permutations, Smouse *et al*. 1986). We considered two types of geographic distances between populations: (i) straight-line Euclidean distance, and (ii) distances measured along road networks, since *S. dioica* often occurs along roads where human activities may facilitate seed dispersal events. Third, population-specific *F_ST_* values were calculated to evaluate the degree of genetic differentiation of each population relative to the others, providing insight into the contribution of each population to the overall genetic structure (Weir and Goudet 2017). Using the R package HIERFSTAT (Goudet 2005), the statistical significance of population-specific *F_ST_* values was assessed through 10,000 bootstrap replicates with 95% confidence intervals. The relationship between population-specific *F*_ST_ and habitat characteristics was then tested using linear regressions including centered and scaled forest and urban habitat surfaces and census population sizes as explanatory variables.

Finally, we used two complementary approaches to investigate the population genetic affiliations at the regional scale. First, a neighbour-joining tree (Saitou and Nei 1987) of populations was built using the *D_A_* genetic distances (Nei *et al*. 1983) and node support was assessed with 1,000 bootstrap replicates using POPTREE2 (Takezaki *et al*. 2010). Second, we performed a Bayesian assignment analysis using STRUCTURE (version 2.3.3, Pritchard *et al*. 2000; Hubisz *et al*. 2009) without prior information on the sampling locations. This method assigns individuals to *K* clusters based on multilocus genotypes, minimizing deviations from Hardy-Weinberg equilibrium and linkage disequilibrium between loci. Ten independent runs for each value of *K* ranging from 1 to 29 (the number of sampled populations) with 1,000,000 Markov chain Monte Carlo (MCMC) iterations and a burn-in period of 100,000 iterations were performed. The most likely value of *K* was inferred by using the *ΔK* statistic, which evaluates the rate of change in log-likelihood between two successive *K* values (Evanno *et al*. 2005).

#### 3. FINE-SCALE PROCESSES: WITHIN-POPULATION SPATIAL GENETIC STRUCTURE AND CONTEMPORARY GENE FLOW

Fine-scale analyses focused on the six intensively sampled populations (100 individuals per population) representing contrasting habitats: three forest and three anthropogenic populations. We evaluated the patterns of fine-scale SGS by testing the relationship between pairwise kinship coefficients (*F_ij_*, Loiselle *et al*. 1995) and pairwise geographic distances between individuals using SPAGeDi (version 1.5, Hardy and Vekemans 2002). Mean *F_ij_* values were computed for eight fixed geographic distance classes (in meters, [0-5[, [5-10[, [10-15[, [15-20[, [20-30[, [30-40[, [40-50[, [60-130[), facilitating visual comparisons among the different populations. The significance of mean *F_ij_* values in each distance class was assessed through 1,000 permutations of individual locations. To compare the strength of the SGS among populations displaying heterogeneity in the spatial distribution of individuals, we calculated the *Sp* statistic, defined as *-b/(1-F^_ij_)*, where *b* is the slope of the regression of *F_ij_* against log-transformed pairwise geographic distances, and *F^_ij_* is the mean kinship between neighbouring individuals (Vekemans and Hardy 2004). The statistical significance of the slopes (*b*) was assessed through 1,000 random permutations of locations among individuals within populations.

In the same six populations, multilocus genotypes of georeferenced adults and collected seeds were then used to perform fractional paternity assignments. This allowed us to estimate the parameters of the pollen dispersal kernel, describing the likelihood of fertilization as a function of distance, and to quantify the proportion of pollen originating from outside the focal population (Klein *et al*. 2006). We applied a spatially explicit mating model that integrates females, males and progenies’ genotypes with spatial coordinates of individuals to estimate, for each seed, the probability of siring by each potential fathers (Oddou-Muratorio *et al*. 2005; Klein *et al*. 2008). This approach allowed us to infer the male reproductive success (*MRS*) and the pollen dispersal distance (*δ*) for each male, as well as the parameters of the population-level pollen dispersal kernel, which was modelled using a negative exponential power function (see Klein *et al*. 2006). Kernel parameters include *b*, a shape parameter describing how sharply siring probability declines with distance, and *µ_δ_*, a scale parameter that reflects the average dispersal distance. Model parameters were estimated with three MCMC chains, with *MRS* and *δ* treated as two latent random variables following Gamma distributions, with means and variances of (*1, σ_MRS_*) and (*µ_δ_, σ_δ_*) respectively. The migration rate (*m*, proportion of seeds for which paternity could not be assigned to a male from the focal population) was also estimated.

Uniform priors were used for *σ_RS_, µ_δ_, σ_δ_, b* and *m* within prior ranges of [0.01,100], [1,100], [0,100], [0.1,10], [0,1] respectively. The convergence of the Markov chains was assessed using three criteria: (i) the Gelman–Rubin diagnostic (< 1), (ii) the Geweke diagnostic (−2 to 2), and (iii) an analysis of chain autocorrelation. We then compared the patterns of pollen flow between forest and anthropogenic habitats using individual dispersal estimates in a generalized linear mixed model, with population included as a random effect.

## RESULTS

### 1. LEVELS OF GENETIC DIVERSITY AT REGIONAL SCALE

Across populations, mean allelic richness (*A_r_*) ranged from 3.0 to 5.2 (Figure 1, Table S2), observed heterozygosity (*H_o_*) ranged from 0.373 to 0.644 and gene diversity (*H_e_*) ranged from 0.424 to 0.719. The mean multilocus intra-population fixation index (*F_IS_*) was 0.057 with 12 out of the 29 populations exhibiting significantly positive *F_IS_* values, indicating departures from random mating (*P* < 0.05). Mapping of *A_r_*, *H_o_*, *H_e_* and *F_IS_* values revealed no obvious geographic gradient or spatial clustering (Figure 1). Neither census population size nor habitat characteristics (i.e., forest and urban habitat surfaces in the one-kilometer radius around each population) affected within-population genetic diversity indices (*A_r_*, *H_o_*, *H_e_*, *F_IS_*, all at *P* > 0.05; Supplementary Table S3). Similarly, the six intensively sampled populations located at opposite ends of the regional habitat spectrum (forest vs. anthropogenic) showed no significant differences in mean levels of genetic diversity or departures from Hardy-Weinberg expectations between habitat types (all at *P >* 0.05).

### 2. LEVELS OF GENETIC DIFFERENTIATION AMONG POPULATIONS AND SPATIAL GENETIC STRUCTURE AT REGIONAL SCALE

Over the whole dataset, we detected significant genetic differentiation, with a multilocus *F_ST_* estimate of 0.069 (*P* < 0.001, 95% confidence interval [0.055, 0.088]). Population pairs exhibited a gradient of genetic differentiation ranging from moderate to strong, with pairwise multilocus *F_ST_* values spanning 0.002 to 0.230. Out of the 378 pairwise *F_ST_* values, 96.6% were significant after Bonferroni correction (Supplementary Table S4). All but one non-significant pairwise *F_ST_* values occurred between populations separated by less than 5 km (Figure 2). A significant pattern of isolation-by-distance was detected, with genetic differentiation between pairwise populations increasing continuously with Euclidean geographic distance (*rz* = 0.210, *P* < 0.001, Figure 2) and the correlation using road-based distances (*rz* = 0.201, *P* < 0.01) was comparable. Population-specific *Fst* values varied among populations (ranging from −0.090 to 0.266) but showed no clear geographic pattern (Figure 1) and were unaffected by either population census size or habitat characteristics (Table S3, all at *P* > 0.05).

**Figure 2.**
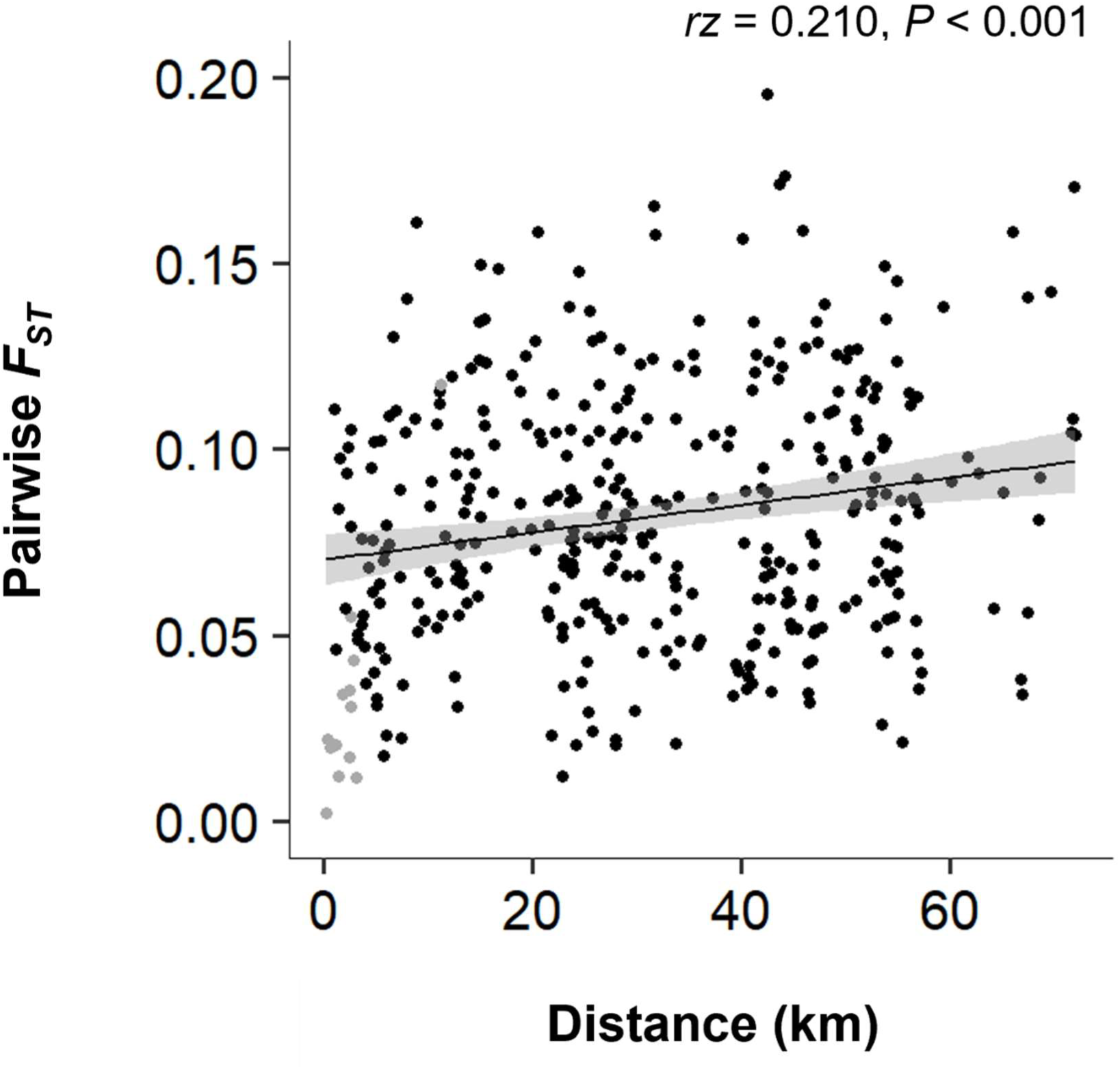
Scatterplot of pairwise *F_ST_* estimates against Euclidean geographical distance in *S. dioica*. Significant pairwise *F_ST_* values are shown in black, and non-significant values in grey.

Genetic affinities among populations were first visualized using a neighbour-joining tree, which revealed a geographically coherent grouping into three distinct sets of populations broadly reflecting the spatial distribution of the sampled populations (Figure 3, A-B). Bayesian assignment analyses identified four genetic clusters (Figure 3, C-D; Supplementary Fig. S2), reflecting, to some extent, the geographic structure depicted in the neighbour-joining tree: populations in the northwest were mostly assigned to a genetically homogeneous cluster, those in the southeast to another cluster, and the intermediate populations showed mixed assignment to the remaining two clusters. However, this geographic signal was heavily blurred by very high levels of admixture, with most individuals exhibiting mixed ancestry across clusters (Figure 3, C).

**Figure 3.**
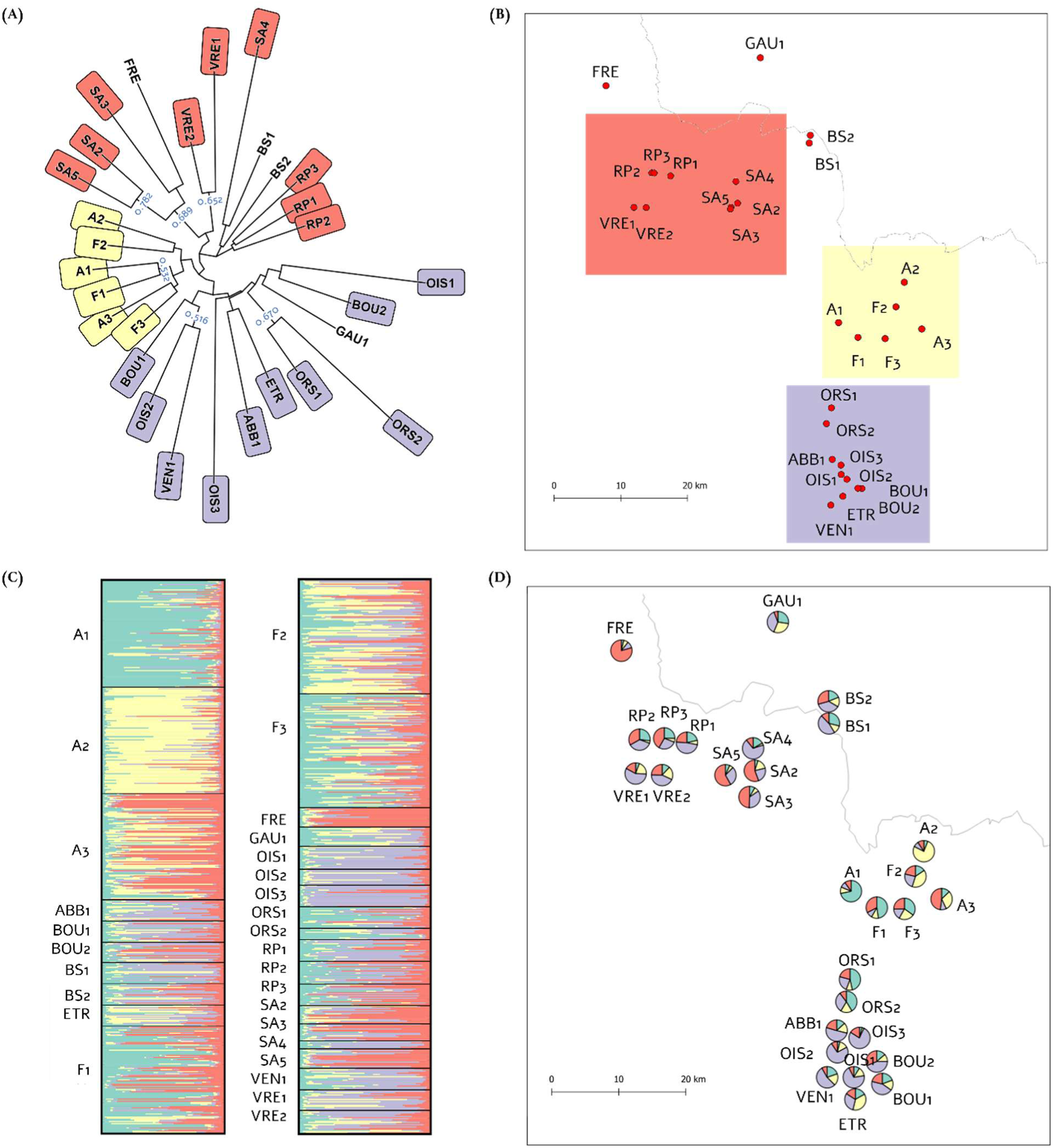
**(A)** Neighbour-joining tree of red campion populations based on Nei’s DA genetic distance (Nei et al., 1983). Blue numbers indicate bootstrap values greater than 50%. **(B)** Sampling locations of populations with colors depicting the geographical distribution of main NJ grouping of populations. **(C)** Bayesian clustering for the most likely number of clusters, *K* = 4. Barplots show individual assignment probabilities to each of the four genetically distinct clusters (each group is represented by a color, each line represents an individual). **(D)** Pie charts representing the mean membership of each population to each of the four most likely clusters.

### 3. FINE-SCALE PROCESSES: WITHIN-POPULATION SPATIAL GENETIC STRUCTURE AND CONTEMPORARY GENE FLOW

A significant pattern of isolation-by-distance was detected within 5 of the 6 extensively surveyed populations, as evidenced by a decreasing level of genetic relatedness with increasing geographical distances between pairs of individuals, and by the *Sp* statistics (Figure 4). At short distance classes (below ten meters) and for all populations, mean kinship coefficients were significantly higher than expected under the null hypothesis of no significant SGS. The *Sp* statistics ranged from 0.004 to 0.022, indicating more than a fivefold variation in the strength of SGS, with the highest values occurring in both forest and anthropogenic habitats (Figure 4). Contemporary pollen dispersal patterns, quantified using paternity analyses, varied among populations, with differences in both the scale and shape of the dispersal kernels, ranging from sharply peaked distributions to a more gradual decline in siring probability with distance (Figure 5, Table 1). Mean pollen dispersal distances within populations were restricted, with values well below 10 m in five of the six populations and reaching 25 m only in the remaining population (Table 1). Analyses of individual-specific pollen dispersal revealed no significant differences between forest and anthropogenic habitats (*F_1,295_* = 0.451*, P* = 0.502), indicating similar dispersal ranges across environments. Notably, these short pollen dispersal distances were coupled with substantial external pollen immigration, with migration rates ranging from 0.35 to 0.53: at least one-third of the pollen cloud in each population thus originated from external donors, irrespective of the habitat context (Table 1).

**Figure 4.**
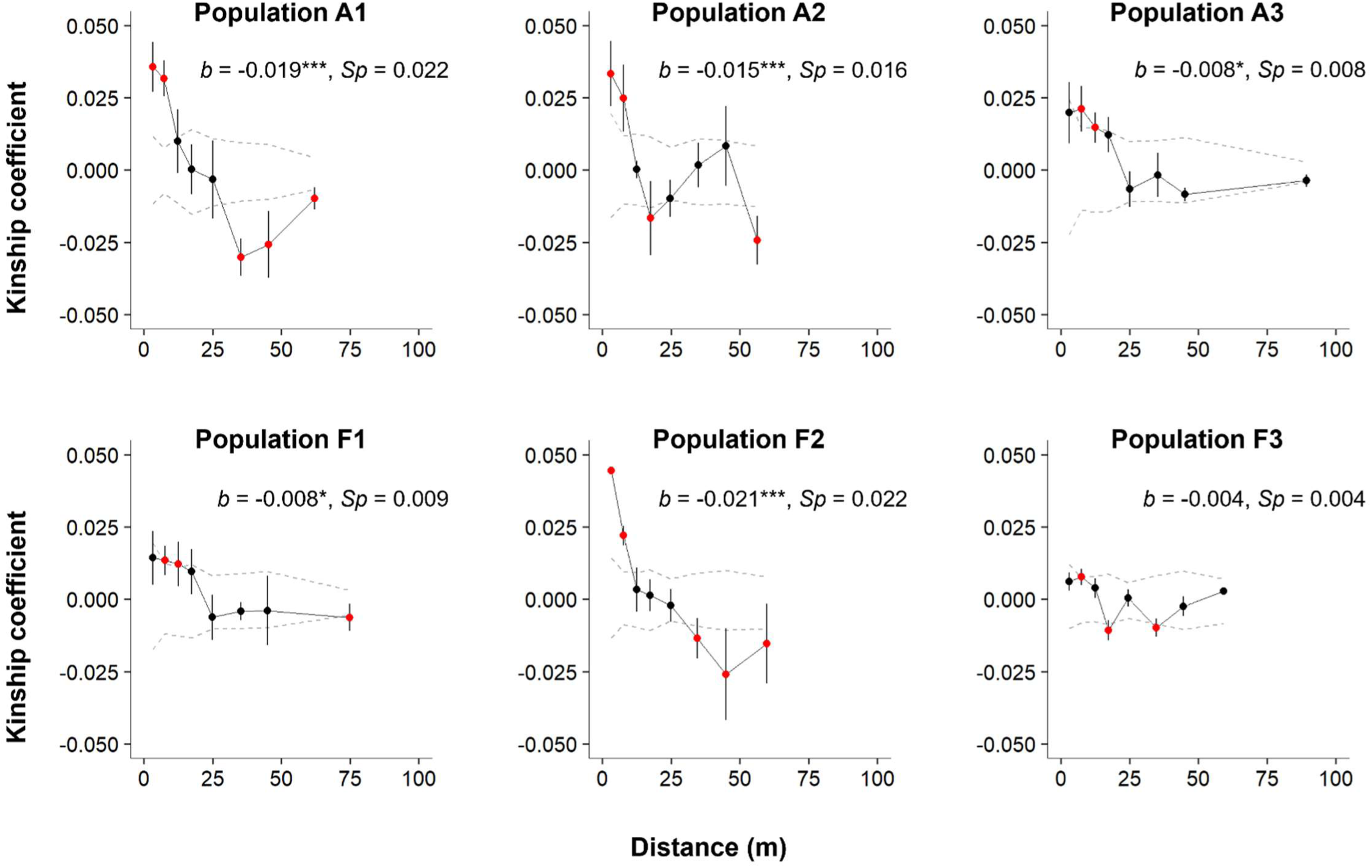
Spatial autocorrelograms showing the relationship between pairwise kinship coefficient (± standard error) calculated following Loiselle et al. (1995) and pairwise geographical distances among individuals (m), for six red campion populations. Populations with names starting with “F” are located in predominantly forested areas, while populations starting with “A” are found in agricultural areas. Standard deviations for each *F_ij_* value per distance class were obtained using jackknifing over loci. Significant mean *F_ij_* values are indicated in red (*P* < 0.05) and significant estimates of *b* (slope of the regression of pairwise *F_ij_* against log-transformed pairwise geographical distances) are indicated by stars (***: *P* < 0.001, *: *P* < 0.05).

**Figure 5.**
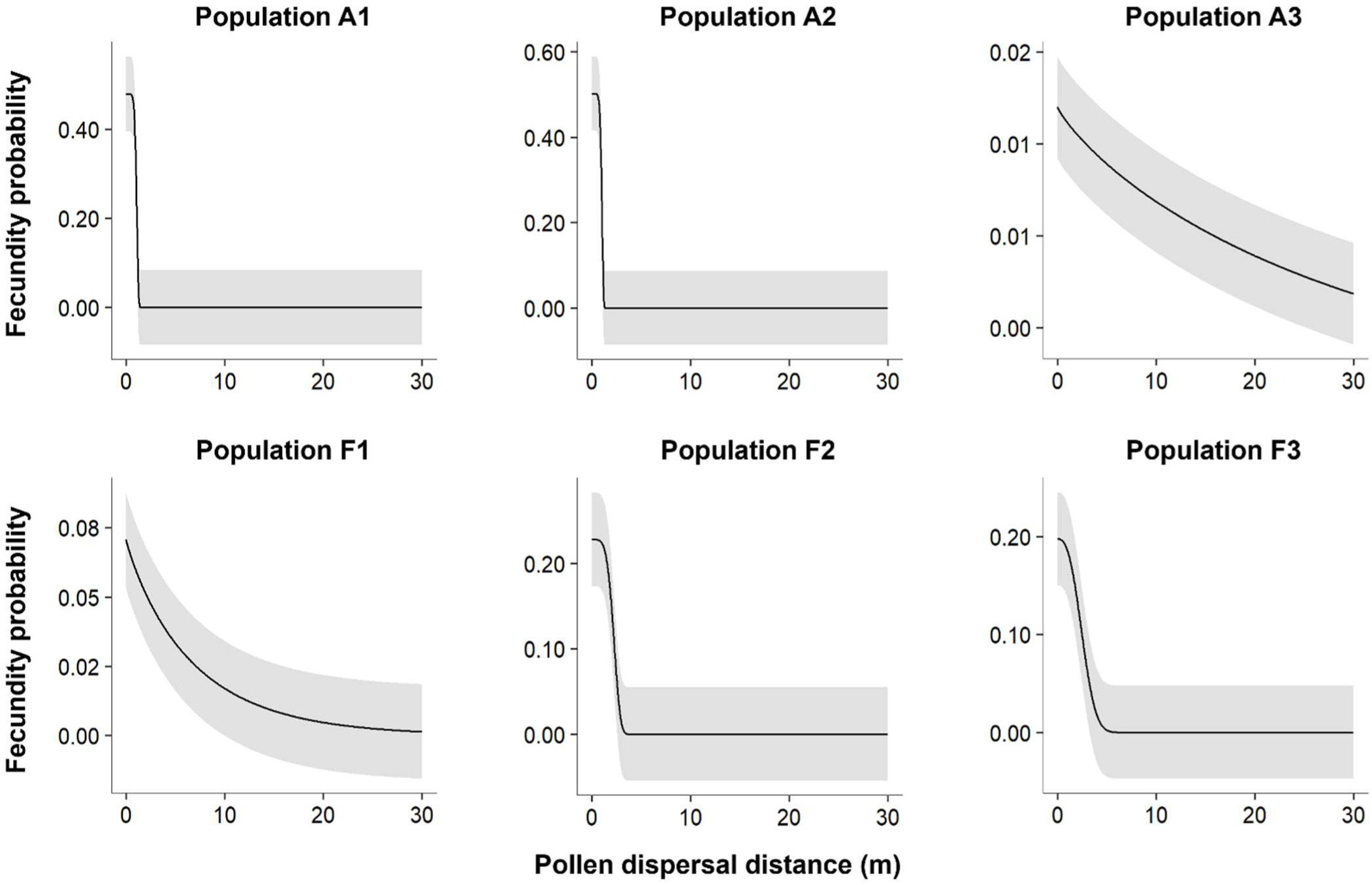
Plots representing the estimated pollen dispersal kernels for six populations of the red campion (*S. dioica*). Populations with names starting with “F” are located in predominantly forested areas, while populations starting with “A” are found in agricultural areas. The grey area represents the standard deviation around the predicted fecundity probability.

**Table 1.** Spatially explicit mating model parameters estimates for six populations of red campion (*S. dioica*). *σ_RS_*: male reproductive success variance; *µ_δ_*: mean pollen dispersal distance (m); *σ_δ_*: variance of pollen dispersal distance; *b*: shape of the pollen dispersal kernel; *m*: migration rate. Populations with names starting with “F” are located in predominantly forested areas, while populations starting with “A” are found in agricultural areas.

| Population | $\sigma_{RS}$ | $\mu_{\delta}$ | $\sigma_{\delta}$ | $b$ | $m$ |
| --- | --- | --- | --- | --- | --- |
| A1 | 0.96 | 2.31 | 4.39 | 7.63 | 0.35 |
| A2 | 1.55 | 2.19 | 4.64 | 8.20 | 0.41 |
| A3 | 1.64 | 25.43 | 2.06 | 0.84 | 0.48 |
| F1 | 2.16 | 6.02 | 10.38 | 0.92 | 0.53 |
| F2 | 1.30 | 4.95 | 2.56 | 4.64 | 0.40 |
| F3 | 1.41 | 5.50 | 3.28 | 2.64 | 0.37 |

## DISCUSSION

Anthropogenic habitat change is classically expected to reduce local genetic diversity and to increase genetic differentiation among plant populations (Young *et al*. 1996; Berge *et al*. 1998; Honnay and Jacquemyn 2007; Aguilar *et al*. 2008; Aavik *et al*. 2013; Alvarado-Serrano *et al*. 2019). Even in common and widespread species, agricultural intensification could modify patterns of SGS, particularly in short-lived species with limited gene dispersal, a characteristic often associated with gravity-dispersed seeds and insect pollination (Van Rossum *et al*. 2004; Auffret *et al*. 2017). Here, we investigated the drivers of neutral genetic variation and its distribution among populations of the red campion (*Silene dioica*) across a land-use gradient. At the regional scale, we observed no detectable effect of habitat type or census population size on levels of genetic diversity and on population genetic differentiation. Moderate but significant population genetic differentiation despite high levels of admixture, as well as a clear isolation-by-distance pattern were detected. Within populations, we found significant fine-scale SGS, consistent with the extremely limited pollen dispersal distances revealed by paternity analyses, but coupled with substantial pollen immigration. Importantly, within-population patterns of SGS and pollen dispersal were not influenced by habitat type. Altogether, our results suggest that gene flow events in this species may be multimodal, combining very short-distance pollen flow within populations with long-distance dispersal of pollen and/or seeds among populations, which likely underpins its resilience to anthropogenic landscape modification. Below, we discuss the main findings of this study in terms of genetic diversity, genetic differentiation, and broad- and fine-scale SGS patterns.

By reducing effective population size and restricting gene flow among previously continuous populations, land-use change can intensify genetic drift and accelerate the erosion of genetic diversity (Ellegren and Galtier 2016; Alvarado-Serrano *et al*. 2019; but see DiLeo *et al*. 2017), with varying outcomes depending on species’ biological traits and specific environmental constraints (Berge *et al*. 1998; Honnay and Jacquemyn 2007; Naaf *et al*. 2022; Guiller *et al*. 2023; see Hardion *et al*. 2026 for strictly urban environments). In our study, we did not detect any effect of anthropization on genetic diversity, in contrast to other studies where negative effects of habitat fragmentation associated with land-use change have been reported (e.g., Aguilar *et al*. 2008; Emel *et al*. 2021; Hardion *et al*. 2026). Several non-mutually exclusive explanations may account for this unexpected result. First, in this common species, populations are typically large, with a mean census size well above 200 across the study area, which likely limits the impact of genetic drift and buffers against the loss of genetic diversity. Second, while studies reporting negative effects of anthropogenic disturbance on genetic diversity often also document concomitant reductions in population size (Alvarado-Serrano *et al*. 2019; Sullivan *et al*. 2019), we detected no association between census population size and habitat characteristics, despite sampling populations across a broad land-use gradient ranging from forest habitats to areas heavily impacted by human activities. Although *S. dioica* is primarily described as a forest-associated species, our results thus suggest that it can maintain large and apparently viable populations even in highly agricultural or urbanized environments, pointing to a certain degree of ecological generalism. Likewise, agricultural intensification is commonly associated with pollinator declines (Ricketts *et al*. 2008; Winfree *et al*. 2009, 2011) and disrupted pollination services (Burd 1994; Ashman *et al*. 2004; Knight *et al*. 2005). However, *S. dioica* is pollinated by a wide range of insect taxa (Kay *et al*. 1984; Barbot *et al*. 2022; Jolivel *et al*. 2026), which may confer resilience to environmental disturbance through functional redundancy among pollinators. Previous studies have reported generally efficient pollination and limited evidence for pollen limitation in this species (e.g., Barbot *et al*. 2022, 2023, but see Kay *et al*. 1984), including in highly agricultural landscapes (Jolivel *et al*. 2026). Such generalist plant-pollinator interactions may therefore prevent recruitment from becoming seed-limited, even under strongly anthropized conditions. Third, the absence of an effect of anthropogenic disturbance on genetic diversity may reflect a temporal lag between human-induced environmental change and the emergence of a detectable genetic response. In other words, several generations are often required before the genetic consequences of a perturbation become detectable, and many more before populations approach a new equilibrium state (Epps and Keyghobadi 2015). *S. dioica* is a short-lived perennial species, with a lifespan of approximately 5-10 years (Giles *et al*. 1998; Goulson 2009). Reproduction is possible from the first year onward, suggesting that generation time is likely on the order of only a few years. The study area has consisted of forest fragments embedded within a conventionally managed agricultural matrix since at least the mid-20th century. Consequently, the populations investigated here have likely been exposed to anthropogenic environmental change over tens generations, suggesting that a genetic signal might have been expected. Moreover, extinction-recolonization dynamics mediated by seed dispersal across the habitat mosaic may continuously reshuffle genetic variation among populations (Whitlock and McCauley 1990; McCauley *et al*. 1995; Auffret *et al*. 2017), while the presence of a persistent seed bank may buffer populations against the loss of genetic diversity associated with environmental change and genetic drift (del Castillo 1994; Honnay *et al*. 2008; Lundemo *et al*. 2009). Beyond metapopulation dynamics and the potential reintroduction of alleles from older populations through seed banks, our results could also be explained by substantial contemporary gene flow among established populations across habitats differing in their degree of anthropogenic alteration. Such exchanges may contribute to maintaining connectivity across the landscape and help explain the regional patterns of SGS observed in this study, as discussed below.

Several lines of evidence indeed support the occurrence of substantial gene flow across the whole study area despite contrasting levels of anthropogenic disturbance. Populations grouped within the same NJ tree clusters were geographically widespread and Bayesian clustering analyses revealed high levels of genetic admixture among the detected population clusters. Furthermore, a significant but moderate level of regional population differentiation was observed in our study (mean multilocus *F_ST_* = 0.069), which is comparable to values typically reported for other outcrossing species pollinated by large insects, such as bumblebees (Gamba and Muchhala 2020). Moderate levels of genetic structure appear to be a common pattern in the red campion and have also been detected in populations from very different environments, including insular archipelagos (Giles and Goudet 1997) and agricultural and forested habitats (Westerbergh and Saura 1994). Finally, paternity analyses indicated high pollen immigration rates within populations. All these results support strong connectivity across the study area and could explain the lack of significant habitat effects on population genetic differentiation. These results are in line with many studies emphasizing that habitat spatial configuration and heterogeneity, rather than proportion of different habitat types alone, are key for pollinator movement (Benton *et al*. 2003; Aavik *et al*. 2013; Kormann *et al*. 2016; Auffret *et al*. 2017; DiLeo *et al*. 2017; Rahimi *et al*. 2021) and for the maintenance of genetic diversity (Naaf *et al*. 2022). In our study region, even in the most intensively farmed areas, the landscape is not a homogeneous arable plain: hedgerows, small forest fragments, and semi-natural features still occur, maintaining potential corridors for pollinator movement.

Our findings are consistent with a metapopulation genetic structure matching a migrant pool model of colonization in which colonizers are drawn at random from multiple sources and where populations follow an island model of population structure once successfully established (Wade and McCauley 1988; Whitlock and McCauley 1990; McCauley *et al*. 1995), a pattern of genetic structure that was also found in a metapopulation network in the red campion in a Swedish archipelago (Giles and Goudet 1997). However, it should be noted that while large-scale dispersal events may occur, we also detected distance-dependent gene flow, as evidenced by a classic isolation-by-distance (IBD) pattern based on Euclidean distances. This IBD pattern reflects a balance between gene flow and genetic drift (Wright 1943; Slatkin 1993; Hutchison and Templeton 1999; Phillipsen *et al*. 2015), whereby gene flow tends to homogenize nearby populations, while increasing geographic distance reduces the likelihood for successful dispersal events and allows genetic drift to amplify genetic differentiation among populations. The higher variance observed in pairwise *F_ST_* values at larger geographic scales suggest that a gene flow/drift equilibrium has not yet been reached in our study region, with genetic drift exerting a stronger influence than gene flow among more distant populations (Hutchison and Templeton 1999; Phillipsen *et al*. 2015). The fact that using road-network distances instead of Euclidean distances did not improve model fit suggests that human activities do not facilitate seed movement in this system, contrary to what has been reported in other systems (Von Der Lippe and Kowarik 2007; Son *et al*. 2024).

In stark contrast with the hypothesis of long-distance dispersal at the regional scale presented above, at finer spatial scales, we detected a significant pattern of IBD within five of the six extensively surveyed populations, suggesting the existence of genetic neighbourhood structures at very fine spatial scales, on the order of 10-20 m. This pattern is consistent with the positive *F_IS_* values observed in most populations (23 of the 29 populations studied, reaching statistical significance in approximately one third of them). Because the red campion is dioecious, the observed fine-scale SGS and local levels of inbreeding cannot be attributed to self-fertilization, but rather to biparental inbreeding owing to spatially restricted pollen and seed dispersal (Heywood 1991; Isagi *et al*. 2007; Lamperty *et al*. 2025). The observed levels of fine-scale SGS are consistent with those reported in other herbaceous insect-pollinated species (e.g., Vekemans and Hardy 2004; Van Rossum and Triest 2007), including populations of *S. dioica* in other regions (Ingvarsson and Giles 1999), and can be linked to the extremely restricted pollen dispersal distances documented with the paternity analyses. Paternity-based studies in herbaceous insect-pollinated species have consistently shown highly leptokurtic pollen dispersal distributions, with most mating events occurring within a few to a few tens of meters (e.g., Smouse *et al*. 1999; Llaurens *et al*. 2008; Van Rossum *et al*. 2011; DiLeo *et al*. 2018; Barbot *et al*. 2025). Short-distance pollen flow in insect pollinated plants likely reflects pollinator foraging behavior driven by optimal foraging strategy (MacArthur and Pianka 1966; Schmitt 1980), combined with limited pollen carry-over capacity. Although *S. dioica* is a generalist species, its main pollinator, *Bombus terrestris*, typically deposits most of its pollen load on the first visited flowers (Joffard *et al*. 2025), promoting short-range movements between neighbouring plants and thereby generating high levels of correlated paternity and fine-scale biparental inbreeding (Hardy *et al*. 2004). Finally, estimated pollen dispersal distances did not differ between populations located in forest and agricultural habitats, suggesting that at short-distance, plant-to-plant movements were comparable across habitat types.

The counterintuitive combination of long-distance pollen flow among populations and restricted within-population dispersal observed in our study species has also been documented in *Pulsatilla vulgaris.* Di Leo *et al*. (2018) indeed found evidence of a multimodal dispersal kernel, where within-population pollination distances were very restricted and shaped by local floral density, while among-population pollen flow was random with respect to population sources. This study also documented a negative effect of the proportion of forest around the focal population on the proportion of immigrant pollen – an effect that was not observed in our study species. The coexistence of short- and long-distance pollinator movements suggests a dual mechanism of pollen transfer: pollinators may enter a population carrying pollen from another, possibly distant, population and subsequently move among neighbouring individuals, thereby introducing alleles from external sources, while maintaining fine-scale SGS.

In conclusion, the lack of significant effects of habitat characteristics on both the levels of genetic diversity and population genetic differentiation supports the occurrence of efficient gene flow over large geographic distances. At the same time, the detection of a significant pattern of isolation-by-distance indicates that geographic distance contributes to shaping spatial genetic structure at the regional scale. Red campion populations appear to conform to a metapopulation functioning, with recurrent recolonization and migration from multiple sources. Finally, our results point to a multimodal pattern of gene flow, combining very short-distance pollen dispersal within each population, regardless of habitat type, with long-distance movement of pollen and/or seeds, which may partly explain the species’ resilience to anthropogenic landscape modification and highlights the remarkable capacity of this widespread herbaceous species to maintain genetic connectivity despite constraints imposed by insect-mediated pollen dispersal and gravity-driven seed dispersal.

## Supplementary Information

Supplementary data are available at *Annals of Botany* online and consist of the following. <u>Table S1</u>: Land-use categories and their classification into four broad habitat types: Forest, Semi-natural, Crop, and Urban. <u>Table S2</u>: Description of 29 sampled populations of the red campion (*S. dioica*). <u>Table S3</u>: Results of the linear regressions examining the relationships between genetic descriptors of the 29 sampled red campions populations, census populations size, and habitat characteristics. <u>Table S4</u>: Pairwise population *F_ST_* values. <u>Fig. S1</u>: PCA results on habitat characteristics within a one-kilometer radius. <u>Fig.</u> <u>S2</u>: Mean *ΔK* statistic (rate of change in log-likelihood between two successive K values) across the successive tested values of *K* (one to 29) over 10 replicates.

## Author contributions

C.J.: Conceptualization; Data curation; Formal analysis; Investigation; Methodology; Project administration; Visualization; Writing – original draft; Writing – review and editing. J-F.A.: Conceptualization; Methodology; Supervision; Validation; Writing – review and editing. E.B.: Conceptualization; Data curation; Formal analysis; Investigation; Methodology; Project administration; Visualization; Writing – review and editing. C.G.: Investigation; Resources. N.J.: Conceptualization; Investigation; Methodology; Supervision; Validation; Writing – review and editing. I.D.C.: Conceptualization; Funding acquisition; Investigation; Methodology; Project administration; Supervision; Validation; Writing – review and editing.

## Acknowledgements

Many thanks to V. Mahut, T. Desort, N. Roscigni and L. Meunier for their help during the field season, as well as P. Tendron, A. Duputié and J. Tonnabel for their contribution to data processing and analysis. This work has been performed using technical support of E. Schmitt, E. Houzé and N. Faure from the Plateforme Serre, cultures et terrains expérimentaux – Université de Lille. We are grateful to L. Debacker and S. Flourez for laboratory assistance. We thank the Service Public de Wallonie as well as the CORINE Biotope project for mapping data. We would like to warmly thank colleagues and collaborators who provided advice, feedback, and discussions on data analyses: P. Milesi, M. Dufaÿ, J. Shykoff, P. David, P. Saumitou-Laprade and M. Monniaux.

