## Supplementary Information for "Local genetic neighbourhoods but resilient gene flow across anthropogenic landscapes in the red campion (*Silene dioica*)"

### Electronic Supplementary Material

**Figure S1.** PCA results on habitat characteristics within a one-kilometer radius. **(A)** Percentage of variance explained by each axis, **(B)** Correlation circle of variables. DistForest: Distance to the closest forest patch (meters); Forest: surface of forest habitat in a one kilometer radius (%); Semi-natural: surface of semi-natural habitat in a one kilometer radius (%); Crop: surface of crops in a one kilometer radius (%); Urban: surface of urban habitat in a one kilometer radius (%).

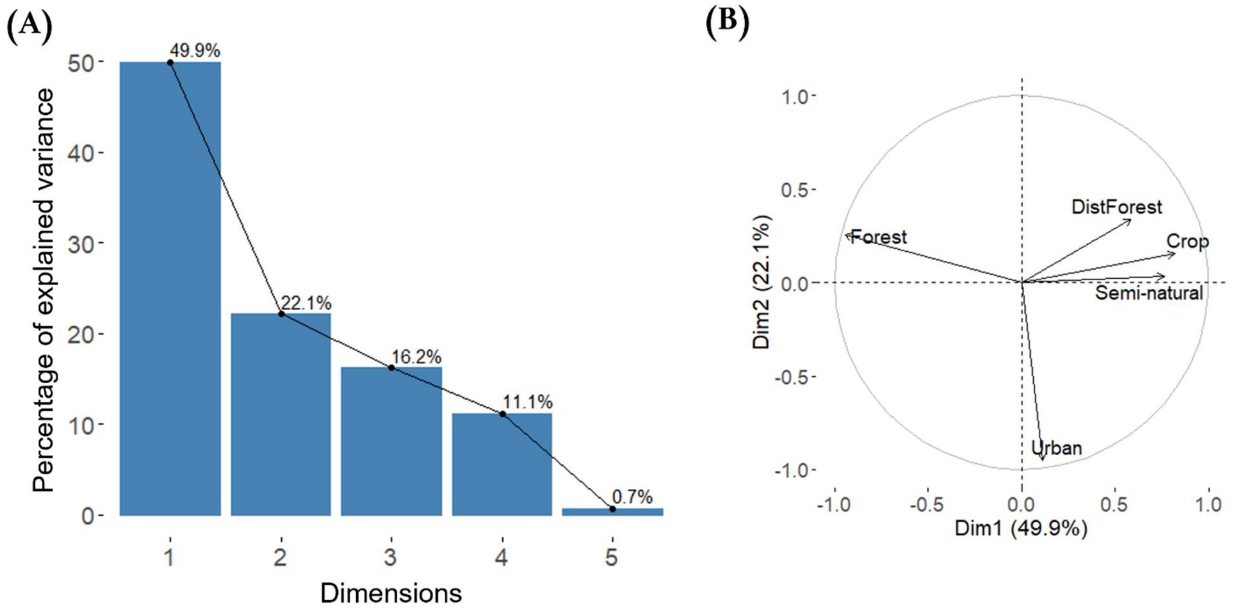

**Figure S2.** Mean *ΔK* statistic (rate of change in log-likelihood between two successive K values) across the successive tested values of *K* (one to 29) over 10 replicates.

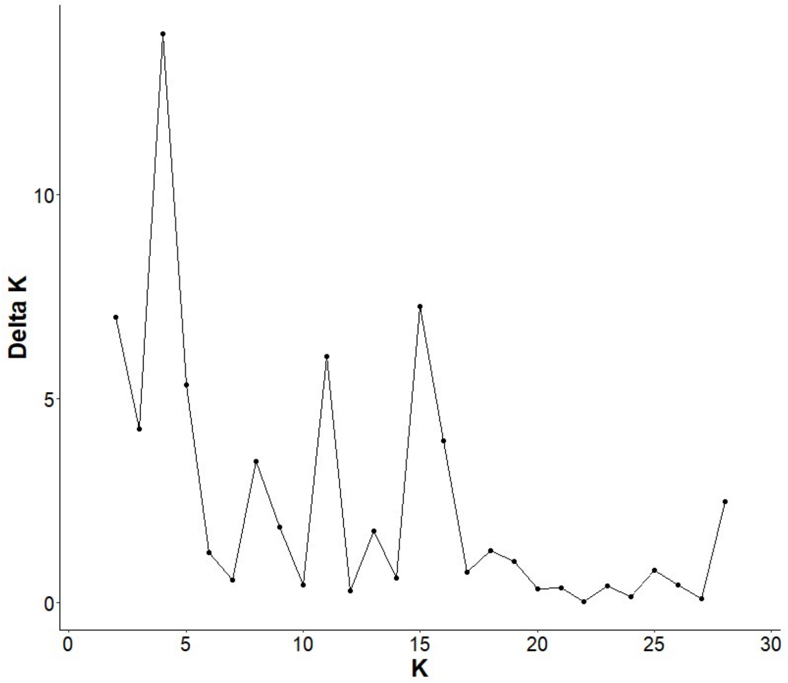

**Table S1.** Land-use categories derived from the four mapping data sources (ARCH, OS Picardie, COSW, and THEIA) and their classification into four broad habitat types: Forest, Semi-natural, Crop, and Urban.

| Habitat type | ARCH | OS Picardie | COSW | THEIA |
|---|---|---|---|---|
| **Forest** | Deciduous forests<br>Conifer plantations<br>Polderian forests<br>Riparian forests | Valley floor deciduous forests<br>Deciduous forests<br>Conifers forests<br>Mixed forests<br>Harvested forests | Deciduous forests<br>Conifers forests<br>Mixed forests<br>Forests | Deciduous forests<br>Coniferous forests |
| **Semi-natural** | Wet meadows<br>Mesophilic meadows<br>Mesophilic pastures<br>Forage meadows<br>Grassy strips<br>Bushes<br>Brownfields<br>Improved meadows<br>Wet tall grass meadows<br>Vegetation belts<br>Submerged vegetation | Natural laws and meadows<br>Meadows<br>Valley floor meadows<br>Moors and bushes | Abandoned lands<br>Brownfields<br>Fallow lands<br>Temporary meadows<br>Permanent meadows | Meadows<br>Grass |
| **Crop** | Orchards<br>Young plantations<br>Poplar plantations<br>Indeterminate plantations<br>Crops | Plots and crops systems<br>Familial gardens<br>Orchards<br>Vineyards<br>Nurseries, market gardening, greenhouses<br>Arable lands | Market gardening<br>Arable lands<br>Crops | Rapeseed<br>Straw cereals<br>Proteaginous<br>Soybeans<br>Sunflower<br>Corn<br>Rice<br>Tubers / Roots<br>Orchards<br>Vineyards |
| **Urban** | Roadsides<br>Urban parks<br>Road networks<br>Towns, villages<br>Industrial sites | Railway infrastructures<br>Roadway infrastructures<br>Urban green spaces<br>Landfills<br>Encampment<br>Cemeteries<br>Airports<br>Building sites<br>Commercial zones<br>Industrial zones<br>Sport equipment<br>Urban fabric<br>Rural habitats<br>Historic habitats<br>Large building complexes | Industrial activities<br>Landfills<br>Active quarries<br>Farm buildings<br>Sport equipment<br>Stores<br>Urban fabric | Dense urban area<br>Sparse urban area<br>Industrial and commercial areas<br>Road |

**Table S2.** Description of 29 sampled populations of the red campion (*S. dioica*). Year: sampling year, Location: municipality in which sampling was conducted; Longitude and Latitude: coordinates in decimal degrees; $N_{census}$: census population size; $N_{sampled}$: number of individuals sampled; DistForest: Distance to the closest forest habitat (meters); Forest: surface of forest habitat in a one kilometer radius (%); SN: surface of semi-natural habitat in a one kilometer radius (%); Crop: surface of crops in a one kilometer radius (%); Urban: surface of urban habitat in a one kilometer radius (%); $A_r$: allelic richness; $H_o$: observed heterozygosity; $H_e$: gene diversity sensus Nei (1978); $F_{IS}$: intra-population fixation index; $F_{ST}$: population-specific $F_{ST}$ according to Weir and Goudet (2017). Significant values of $F_{IS}$ fixation index and $F_{ST}$ are indicated by stars (***: $P < 0.001$, **: $P < 0.01$, *: $P < 0.05$). *P:* P-value.

Nei M. 1978. Estimation of average heterozygosity and genetic distance from a small number of individuals. Genetics 89: 583–590.

Weir BS, Goudet J. 2017. A unified characterization of population structure and relatedness. Genetics 206: 2085–2103.

| ID | Year | Location | Longitude | Latitude | $N_{census}$ | $N_{sampled}$ | DistForet | Forest | SN | Crop | Urban | $A_r$ | $H_o$ | $H_e$ | $F_{IS}$ | $F_{ST}$ |
|---|---|---|---|---|---|---|---|---|---|---|---|---|---|---|---|---|
| **A1** | 2023 | Potelle | 3.6598483 | 50.2326281 | 800 | 100 | 0 | 0.07 | 0.44 | 0.37 | 0.11 | 4.7 | 0.610 | 0.679 | **0.102*****| -0.035 |
| **A2** | 2023 | Bavay | 3.7996608 | 50.2868433 | 2000 | 100 | 0 | 0.15 | 0.31 | 0.41 | 0.13 | 4.2 | 0.504 | 0.559 | **0.098***** | **0.148*** |
| **A3** | 2023 | Pont-sur-Sambre | 3.8355342 | 50.2229467 | 300 | 100 | 71.00 | 0.09 | 0.47 | 0.31 | 0.12 | 5.1 | 0.644 | 0.656 | 0.018 | 0.000 |
| **ABB1** | 2019 | Rejet-de-Beaulieu | 3.6441633 | 50.0466608 | 138 | 20 | 0 | 0.05 | 0.57 | 0.31 | 0.04 | 4.5 | 0.560 | 0.524 | -0.068 | 0.199 |
| **BOU1** | 2019 | Boué | 3.7062825 | 50.0066967 | 625 | 20 | 0 | 0.60 | 0.15 | 0.05 | 0.17 | 4.8 | 0.490 | 0.613 | **0.201***** | **0.070*** |
| **BOU2** | 2019 | Boué | 3.6976417 | 50.007015 | 39 | 19 | 0 | 0.39 | 0.23 | 0.05 | 0.31 | 4.9 | 0.546 | 0.622 | **0.123*** | 0.054 |
| **BS1** | 2019 | Condé-sur-l'Escaut | 3.6008908 | 50.477405 | 93 | 20 | 0 | 0.65 | 0.16 | 0.09 | 0.11 | 5.2 | 0.630 | 0.622 | -0.014 | 0.052 |
| **BS2** | 2019 | Condé-sur-l'Escaut | 3.6033158 | 50.4880025 | 46 | 20 | 0 | 0.72 | 0.02 | 0.11 | 0.14 | 4.4 | 0.540 | 0.581 | 0.071 | **0.115*** |
| **ETR** | 2019 | Etreux | 3.665685 | 49.996355 | 54 | 20 | 226.53 | 0.03 | 0.23 | 0.48 | 0.23 | 4.4 | 0.486 | 0.602 | **0.193**** | **0.087*** |
| **F1** | 2023 | Locquignol | 3.7006172 | 50.2124031 | 150 | 100 | 0 | 0.98 | 0.02 | 0.00 | 0.00 | 4.8 | 0.493 | 0.557 | **0.114***** | **0.151*** |
| **F2** | 2023 | Locquignol | 3.7815922 | 50.2535217 | 250 | 100 | 0 | 0.80 | 0.18 | 0.00 | 0.02 | 4.5 | 0.562 | 0.584 | 0.037 | 0.110 |
| **F3** | 2023 | Locquignol | 3.7580894 | 50.2103733 | 400 | 100 | 0 | 0.77 | 0.17 | 0.04 | 0.01 | 5.2 | 0.640 | 0.651 | 0.017 | 0.008 |
| **FRE** | 2018 | Templeuve-en-Pévèle | 3.1701685 | 50.5567963 | 42 | 17 | 0 | 0.13 | 0.34 | 0.35 | 0.15 | 3.1 | 0.588 | 0.562 | -0.046 | **0.142*** |
| **GAU1** | 2019 | Tournai (Belgium) | 3.498345 | 50.5940175 | 178 | 17 | 0 | 0.16 | 0.19 | 0.11 | 0.53 | 3.8 | 0.588 | 0.560 | -0.051 | **0.145*** |
| **OIS1** | 2019 | Oisy | 3.6626792 | 50.02587 | 148 | 20 | 1,268.89 | 0.00 | 0.34 | 0.56 | 0.10 | 4.2 | 0.642 | 0.645 | 0.005 | 0.017 |
| **OIS2** | 2019 | Oisy | 3.674575 | 50.0194143 | 25 | 14 | 412.23 | 0.03 | 0.55 | 0.27 | 0.11 | 3.8 | 0.557 | 0.513 | -0.087 | **0.216*** |
| **OIS3** | 2019 | Fesmy-le-Sart | 3.6623211 | 50.0387 | 59 | 19 | 173.11 | 0.03 | 0.66 | 0.22 | 0.05 | 5 | 0.526 | 0.686 | **0.233***** | -0.037 |
| **ORS1** | 2019 | Ors | 3.6433133 | 50.1168417 | 618 | 19 | 0 | 0.44 | 0.31 | 0.2. | 0.04 | 4.7 | 0.600 | 0.622 | 0.035 | 0.053 |
| **ORS2** | 2019 | Ors | 3.63263 | 50.0952633 | 10 | 10 | 0 | 0.02 | 0.60 | 0.15 | 0.19 | 3 | 0.373 | 0.488 | **0.233*** | 0.266 |
| **RP1** | 2019 | Marchiennes | 3.3065456 | 50.4338289 | 121 | 19 | 0 | 0.53 | 0.08 | 0.32 | 0.07 | 4.7 | 0.589 | 0.611 | 0.035 | 0.069 |
| **RP2** | 2018 | Beuvry-la-Forêt | 3.2667383 | 50.438065 | 176 | 20 | 0 | 0.51 | 0.10 | 0.27 | 0.12 | 4.6 | 0.628 | 0.615 | -0.021 | 0.062 |
| **RP3** | 2019 | Beuvry-la-Forêt | 3.2720429 | 50.437919 | 179 | 19 | 0 | 0.51 | 0.10 | 0.27 | 0.12 | 4.6 | 0.554 | 0.592 | 0.064 | 0.099 |
| **SA2** | 2019 | Raismes | 3.4482052 | 50.3960708 | 31 | 16 | 0 | 0.53 | 0.08 | 0.10 | 0.12 | 4.7 | 0.482 | 0.584 | **0.174**** | 0.114 |
| **SA3** | 2018 | Raismes | 3.4341156 | 50.3906056 | 21 | 15 | 0 | 0.36 | 0.09 | 0.16 | 0.11 | 4.8 | 0.568 | 0.608 | 0.066 | 0.076 |
| **SA4** | 2019 | Saint-Amand-les-Eaux | 3.4449333 | 50.4259333 | 7 | 7 | 0 | 0.71 | 0.12 | 0.03 | 0.14 | 3.2 | 0.543 | 0.424 | **0.281*** | *NA* |
| **SA5** | 2019 | Raismes | 3.4330632 | 50.38825 | 113 | 17 | 0 | 0.30 | 0.09 | 0.20 | 0.16 | 4.6 | 0.412 | 0.523 | **0.212***** | **0.209*** |
| **VEN1** | 2019 | Vénérolles | 3.6402917 | 49.9844033 | 69 | 19 | 0 | 0.02 | 0.26 | 0.60 | 0.10 | 3.7 | 0.635 | 0.637 | 0.003 | 0.028 |
| **VRE1** | 2018 | Vred | 3.2288759 | 50.3910685 | 29 | 18 | 0 | 0.14 | 0.30 | 0.26 | 0.26 | 4.7 | 0.604 | 0.719 | **0.160*** | -0.090 |
| **VRE2** | 2018 | Rieulay | 3.2546317 | 50.3907883 | 88 | 20 | 0 | 0.20 | 0.26 | 0.36 | 0.06 | 4.2 | 0.600 | 0.608 | 0.013 | 0.073 |

**Table S3.** Results of the linear regressions examining the relationships between genetic descriptors of the 29 sampled red campions populations ($A_r$: allelic richness; $H_o$: observed heterozygosity; $H_e$: gene diversity sensus Nei (1978); $F_{IS}$: intra-population fixation index; $F_{ST}$: population-specific $F_{ST}$ according to Weir and Goudet (2017), census populations size, and habitat characteristics (percentage of forest and urban cover within a one kilometer radius). All predictors were centered and scaled (mean = 0, SD = 1).

| Response variable | Predictor | Estimate | *P* |
|---|---|---|---|
| $A_r$ | Forest habitat | 0.174 | 0.142 |
| | Urban habitat | -0.098 | 0.403 |
| | Census population size | 0.070 | 0.531 |
| $H_o$ | Forest habitat | -0.005 | 0.707 |
| | Urban habitat | -0.010 | 0.501 |
| | Census population size | -0.003 | 0.824 |
| $H_e$ | Forest habitat | -0.011 | 0.400 |
| | Urban habitat | -0.007 | 0.602 |
| | Census population size | 0.004 | 0.756 |
| $F_{IS}$ | Forest habitat | -0.012 | 0.620 |
| | Urban habitat | 0.005 | 0.844 |
| | Census population size | 0.013 | 0.572 |
| $F_{ST}$ | Forest habitat | 0.004 | 0.840 |
| | Urban habitat | 0.007 | 0.701 |
| | Census population size | -0.002 | 0.906 |

42 **Table S4.** Pairwise population $F_{ST}$ values. The color gradient reflects the range of $F_{ST}$ values, with
43 grey cells indicating non-significant comparisons ($P < 0.05$ after Bonferroni corrections).

| | A1 | A2 | A3 | ABB1 | BOU1 | BOU2 | BS1 | BS2 | ETR | F1 | F2 | F3 | FRE | GAU1 | OIS1 | OIS2 | OIS3 | ORS1 | ORS2 | RP1 | RP2 | RP3 | SA2 | SA3 | SA5 | VEN1 | VRE1 |
|---|---|---|---|---|---|---|---|---|---|---|---|---|---|---|---|---|---|---|---|---|---|---|---|---|---|---|---|
| **A2** | 0.077 | | | | | | | | | | | | | | | | | | | | | | | | | | |
| **A3** | 0.039 | 0.037 | | | | | | | | | | | | | | | | | | | | | | | | | |
| **ABB1** | 0.104 | 0.083 | 0.078 | | | | | | | | | | | | | | | | | | | | | | | | |
| **BOU1** | 0.029 | 0.053 | 0.024 | 0.074 | | | | | | | | | | | | | | | | | | | | | | | |
| **BOU2** | 0.043 | 0.086 | 0.059 | 0.044 | 0.020 | | | | | | | | | | | | | | | | | | | | | | |
| **BS1** | 0.077 | 0.137 | 0.085 | 0.139 | 0.093 | 0.114 | | | | | | | | | | | | | | | | | | | | | |
| **BS2** | 0.054 | 0.105 | 0.063 | 0.126 | 0.045 | 0.075 | 0.046 | | | | | | | | | | | | | | | | | | | | |
| **F1** | 0.056 | 0.021 | 0.022 | 0.070 | 0.012 | 0.055 | 0.103 | 0.077 | 0.087 | | | | | | | | | | | | | | | | | | |
| **F2** | 0.053 | 0.107 | 0.054 | 0.086 | 0.052 | 0.050 | 0.123 | 0.066 | 0.030 | 0.089 | | | | | | | | | | | | | | | | | |
| **F3** | 0.051 | 0.047 | 0.033 | 0.112 | 0.021 | 0.071 | 0.103 | 0.046 | 0.037 | 0.037 | 0.031 | | | | | | | | | | | | | | | | |
| **FRE** | 0.022 | 0.059 | 0.018 | 0.079 | 0.012 | 0.036 | 0.071 | 0.166 | 0.105 | 0.149 | 0.124 | 0.092 | | | | | | | | | | | | | | | |
| **GAU1** | 0.124 | 0.135 | 0.091 | 0.159 | 0.104 | 0.108 | 0.158 | 0.089 | 0.056 | 0.052 | 0.067 | 0.035 | 0.138 | | | | | | | | | | | | | | |
| **OIS1** | 0.052 | 0.075 | 0.052 | 0.098 | 0.034 | 0.038 | 0.134 | 0.116 | 0.049 | 0.102 | 0.083 | 0.055 | 0.093 | 0.057 | | | | | | | | | | | | | |
| **OIS2** | 0.069 | 0.075 | 0.058 | 0.105 | 0.056 | 0.050 | 0.127 | 0.098 | 0.079 | 0.080 | 0.085 | 0.063 | 0.142 | 0.088 | 0.111 | | | | | | | | | | | | |
| **OIS3** | 0.089 | 0.108 | 0.076 | 0.076 | 0.031 | 0.057 | 0.127 | 0.095 | 0.076 | 0.107 | 0.102 | 0.073 | 0.141 | 0.094 | 0.084 | 0.101 | | | | | | | | | | | |
| **ORS1** | 0.086 | 0.116 | 0.076 | 0.098 | 0.062 | 0.068 | 0.111 | 0.121 | 0.083 | 0.056 | 0.120 | 0.067 | 0.138 | 0.054 | 0.067 | 0.112 | 0.108 | | | | | | | | | | |
| **ORS2** | 0.055 | 0.115 | 0.078 | 0.104 | 0.067 | 0.031 | 0.157 | 0.171 | 0.117 | 0.099 | 0.159 | 0.123 | 0.230 | 0.112 | 0.141 | 0.161 | 0.130 | 0.035 | | | | | | | | | |
| **RP1** | 0.107 | 0.148 | 0.129 | 0.102 | 0.116 | 0.064 | 0.217 | 0.023 | 0.074 | 0.087 | 0.034 | 0.039 | 0.149 | 0.088 | 0.119 | 0.052 | 0.083 | 0.124 | 0.174 | | | | | | | | |
| **RP2** | 0.042 | 0.101 | 0.059 | 0.116 | 0.021 | 0.061 | 0.057 | 0.054 | 0.086 | 0.041 | 0.090 | 0.046 | 0.124 | 0.068 | 0.088 | 0.067 | 0.088 | 0.053 | 0.127 | 0.043 | | | | | | | |
| **RP3** | 0.049 | 0.125 | 0.069 | 0.105 | 0.040 | 0.045 | 0.073 | 0.021 | 0.087 | 0.042 | 0.060 | 0.035 | 0.150 | 0.076 | 0.101 | 0.074 | 0.085 | 0.101 | 0.159 | 0.017 | 0.022 | | | | | | |
| **SA2** | 0.047 | 0.116 | 0.058 | 0.108 | 0.036 | 0.054 | 0.067 | 0.082 | 0.075 | 0.096 | 0.076 | 0.066 | 0.130 | 0.104 | 0.122 | 0.068 | 0.073 | 0.122 | 0.135 | 0.052 | 0.087 | 0.064 | | | | | |
| **SA3** | 0.086 | 0.090 | 0.065 | 0.048 | 0.051 | 0.044 | 0.122 | 0.088 | 0.060 | 0.052 | 0.088 | 0.046 | 0.075 | 0.071 | 0.070 | 0.059 | 0.070 | 0.069 | 0.101 | 0.085 | 0.065 | 0.065 | 0.021 | | | | |
| **SA5** | 0.070 | 0.079 | 0.049 | 0.037 | 0.051 | 0.032 | 0.111 | 0.101 | 0.109 | 0.067 | 0.113 | 0.076 | 0.117 | 0.098 | 0.119 | 0.062 | 0.095 | 0.108 | 0.121 | 0.091 | 0.075 | 0.069 | 0.012 | 0.002 | | | |
| **VEN1** | 0.105 | 0.127 | 0.087 | 0.042 | 0.077 | 0.043 | 0.135 | 0.115 | 0.093 | 0.129 | 0.124 | 0.077 | 0.170 | 0.081 | 0.102 | 0.095 | 0.109 | 0.061 | 0.120 | 0.086 | 0.083 | 0.114 | 0.097 | 0.101 | 0.134 | | |
| **VRE1** | 0.068 | 0.126 | 0.104 | 0.110 | 0.059 | 0.040 | 0.145 | 0.105 | 0.067 | 0.105 | 0.066 | 0.060 | 0.116 | 0.085 | 0.060 | 0.097 | 0.058 | 0.088 | 0.129 | 0.066 | 0.073 | 0.080 | 0.068 | 0.075 | 0.093 | 0.102 | |
| **VRE2** | 0.061 | 0.084 | 0.051 | 0.110 | 0.055 | 0.065 | 0.111 | 0.055 | 0.065 | 0.104 | 0.036 | 0.048 | 0.125 | 0.092 | 0.097 | 0.085 | 0.093 | 0.134 | 0.196 | 0.023 | 0.064 | 0.047 | 0.059 | 0.093 | 0.099 | 0.117 | 0.034 |

| | |
|---|---|
| (grey) | $F_{ST}$ NS |
| (white) | $F_{ST} < 0.05$ |
| (pale yellow) | $0.05 < F_{ST} < 0.10$ |
| (yellow) | $0.10 < F_{ST} < 0.15$ |
| (dark gold) | $0.15 < F_{ST} < 0.20$ |
| (orange) | $F_{ST} > 0.20$ |

44
